# pyreml: a general-purpose, GPU-accelerated mixed model solver in Python

**DOI:** 10.64898/2026.10.20.757556

**Authors:** Alexandre Marchal, Catherine Trottier, Xavier Bry, Daniele Raimondi

**Affiliations:** Institut de Génétique Moléculaire de Montpellier (IGMM), UMR 5535, Université de Montpellier, CNRS, Montpellier, France; Institut Montpelliérain Alexander Grothendieck (IMAG), UMR 5149, Université de Montpellier, CNRS, Montpellier, France

**Keywords:** Prediction, Linear mixed model, GPU, REML, GBLUP

## Abstract

Linear mixed models (LMM) are a popular modeling and inference framework, widely used in bioinformatics and other scientific fields. To obtain an unbiased estimation of the variance components, an LMM has to be fitted using restricted maximum likelihood (REML), a computationally intensive step. Most REML software is closed source, and currently there is no `Python` library offering complete LMM functionalities, such as the specification of spatial and genomic variance structures, multivariate analyses, random regression and factors, and heteroscedastic residuals. Here we propose `pyreml`, an LMM solver that fills this gap, comes with a user-centered interface and speeds up calculations by running on GPU. We show that `pyreml` fits a genomic best linear unbiased prediction (GBLUP) model more than one order of magnitude (*>* 12 ×) faster than the current state-of-the-art, closed source REML software. We also show that `pyreml` can estimate a latent factor-analytic variance structure, a challenging feature that no current open source REML solution implements. Finally, we illustrate the use of `pyreml` to analyze longitudinal data, and to perform spatial analysis. Anchored by 500+ functional tests assessing the equivalence of its outputs to those from reference software (`lme4`, `rrBLUP` and `BLUPF90`), we demonstrate `pyreml` to be a performant, reliable and open-source solution available for academic, research and industrial LMM users.

## 1 Introduction

Notwithstanding the increasing ubiquity of complex non-linear models in any application of data science, Linear Mixed Models (LMMs) remain a major inference solution in many scientific fields such as genetics [Crossa et al., 2025, Cooper et al., 2025], spatial statistics [Lark et al., 2006, De Caires et al., 2025], ecology [Oberpriller et al., 2022], and health science [Ouyang et al., 2024, Karimi Ghahfarokhi et al., 2025].

LMMs assume that some effects are random, and moderate their predictions in proportion to the uncertainty attached to them. The less evidence supports a given level, the closer to the population mean it is predicted. The intuition is familiar: a speaker who tempers claims resting on thin evidence earns more credit than one asserting everything with equal confidence. Machine learning applies the same principle under the name of regularization. LMMs apply it analytically, which is what makes their predictions generalize better beyond the training data [Henderson, 1975]. It also allows the uncertainty itself to be structured, for instance by genomic relatedness or spatial proximity. All of this rests on variance parameters, estimated by restricted maximum likelihood (REML) [Patterson and Thompson, 1971, Harville, 1977], a computationally intensive step.

Historically, REML software is dominated by closed-source binaries with non-commercial licenses (`Wombat` [Meyer, 2006], `BLUPF90` [Misztal et al., 2014]), by subscription-based solutions (`ASReml` [Butler et al., 2009], `SAS` [SAS Institute Inc., 2023]), or by free solutions with severe functional limitations: `lme4` [Bates et al., 2015], `rrBLUP` [Endelman, 2011], `glimix` [Lippert et al., 2014], `spaMM` [Rousset and Ferdy, 2014]. As a notable counter-example, `sommer` [Giovanny, 2016] proposes a rich, open implementation targeting `R` users. Even though Python is the most popular language for data science and bioinformatics, it currently lacks a flexible, modern and free LMM library that could compete with proprietary software in terms of functionalities and performance.

Here we propose `pyreml`, a `PyTorch`-based [Paszke et al., 2019] general purpose LMM solver, which is the first library performing REML estimations on GPU. It also runs on CPU, addressing use cases ranging from small experimental designs to heavy industrial routines.

The combination of mathematical and GPU optimizations that we propose allows `pyreml` to process data faster than state-of-the-art methods, improving the scalability of LMMs to datasets of any size, while still performing exact computation.

## 2 Methods

### 2.1 How pyreml works

`pyreml` fits the LMM: **y** = **X***β* + **Zu** + **r** with **u** ∼ N (**0, G**) and **r** ∼ N (**0, R**), with **y** being the response (label), *β* and **u** respectively the fixed and random effects, and **X**_*n*×*p*_ and **Z**_*n*×*q*_ incidence matrices (features) (see Suppl. S1). LMM solvers classically use the expectation-maximization (EM) [Dempster et al., 1977] and average-information (AI) [Gilmour et al., 1995] algorithms, sometimes accelerated by parameter expansion (PX) [Liu et al., 1998, Meyer, 2006, Butler et al., 2009]. On the other hand, Simchoni and Rosset [2023] suggested direct Gradient Descent (GD) of the log-likelihood. `pyreml` uses GD but extended the work of Simchoni and Rosset [2023] by adding the REML corrective term to their loss, to tackle the bias of variance estimates [Harville, 1977] (see Suppl. S1.3.1).

`pyreml` is based on `PyTorch` [Paszke et al., 2019], and it can be run either on CPU or on GPU, at user discretion. We use a limited-memory L-BFGS [Liu and Nocedal, 1989] optimizer using a strong Wolfe line search, and an Adaptive Moment Estimation (Adam) [Kingma and Ba, 2017] fallback to handle occasionally occurring singularity exceptions (see Suppl. S1.3.2). Convergence is declared when reaching a dynamic threshold that estimates the number of significant digits of the likelihood (see Suppl. S1.3.6). `pyreml` offers 3 strategies for the REML step: i) running with double precision (FP64), ii) with float32 (FP32), and iii) a mixed precision heuristic to minimize the overhead that FP64 computation carries on consumer GPUs, while limiting the risk of numeric instabilities that can derail a pure FP32 optimization (see Suppl. S1.1). When *q < n*, `pyreml` automatically uses the Sherman-Morrison-Woodbury (SMW) identity to reduce the dimensionality of the problem (see Suppl. S1.3.3). `pyreml` exploits the factored form of each variance component to assemble the total variance without ever forming its Kronecker blocs nor **ZGZ**^⊤^ (see Suppl. S1.3.4), and replaces automatic differentiation through the Cholesky factorisation by the closed-form REML gradient (see Suppl. S1.3.5). Moreover, the representation of large matrices such as **Z** or **R** has been replaced by index grids wherever the algebra allows, which strongly reduces their (V)RAM footprint.

### 2.2 How to use pyreml

`pyreml` is distributed as an open-source python library with an extensive documentation and a user-centered, high-level API. It takes a `pandas` [McKinney, 2010] dataframe as input and internally manages the missing data. It reads a `patsy` [Smith and patsy contributors, 2024] formula for the fixed effects, can predict a single or multiple real-valued targets (multivariate analysis) and another `patsy` formula for each random effect (random regression, random factors) in a way that is similar to the `lme4` API.

`pyreml`’s core design choice is to define every random effect and the residual as a Kronecker product: **G** = ⨁**_a_ Σ_a_** ⊗ **K_a_**. This way, both the left-hand term **Σ_a_** and right-hand term **K_a_** of each random effect and of the residual can be specified. The overall philosophy is to provide a flexible and unified framework for a large variety of applications. A total of 8 covariance structures on the left-hand term (i.i.d., diagonal, unstructured, factor-analytic, 2 bloc-diagonal and 2 proportional-bloc forms) and 7 right-hand structure across levels (identity, known matrix, heteroscedastic, and 4 decay kernels) can be combined. Exhaustive specifications for **K_a_** and **Σ_a_** are detailed in the package documentation.

As notable applications, **K_a_** may be a genomic kinship matrix, *i*.*e*. the genomic best linear unbiased prediction (GBLUP) model [Meuwissen et al., 2001, VanRaden, 2008], or a matrix of spatial distance [Lark et al., 2006]. It can also encode heteroscedasticity modeled as **K_a_** = diag (exp(*V* **h**)), with *V* an heteroscedasticity incidence matrix specified with a `patsy` formula and **h** a vector of parameters learnt from the data.

On the other hand, **Σ_a_** gathers all the actual variances and covariances to estimate: for instance between different responses. It may be approximated with a reduced rank factor analytic (FA) model [Smith et al., 2001, Kirkpatrick and Meyer, 2004]: **Σ_a_** = **Q_a_Λ_a_Q_a_**^⊤^ + **Ψ_a_**, with **Q_a_Λ_a_Q_a_**^⊤^ a principal component decomposition. A `jitter` argument allows a simultaneously absolute and relative shrinkage of the covariance as to defuse singularities met in the gradient descent process 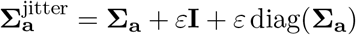, with *ε* the jitter.

### 2.3 Functional tests

The `pyreml` repository ships a suite of functional tests asserting either numerical agreement with established mixed model software (`lme4`, `rrBLUP`, `BLUPF90`), or absolute performance (see Suppl. S2). Each reference is generated once by the competing software and frozen as a JSON file versioned alongside the code. As is standard software practice, the suite runs automatically in the continuous integration pipeline. A total of 505 tests pass, and 6 assertions are marked as expected failures due to lack of identifiability on specific edge cases, as documented in Suppl. S2. Using FP32 computation breaks 241 of the functional tests, therefore this option is never advisable.

### 2.4 Hardware

The benchmarks and applications were conducted on 2 platforms: (i) a server with an AMD EPYC 9124 CPU (2×16 physical cores, 64 logical threads, thereafter simply referred as “64 CPUs”) and a NVIDIA L40S GPU, (ii) a cloud instance with a NVIDIA A100 (PCIe) GPU.

## 3 Results

### 3.1 pyreml brings large-scale REML to the GPU

In Fig. 1a, we benchmarked `pyreml` scalability against `Wombat` [Meyer, 2006], which together with the commercially licensed `ASReml` [Butler et al., 2009] are the *de facto* standard for largescale REML in quantitative genetics. `Wombat` was run on 64 CPUs using EM, AI and PX-AI (a mix of PX and AI steps) algorithms.

**Figure 1.**
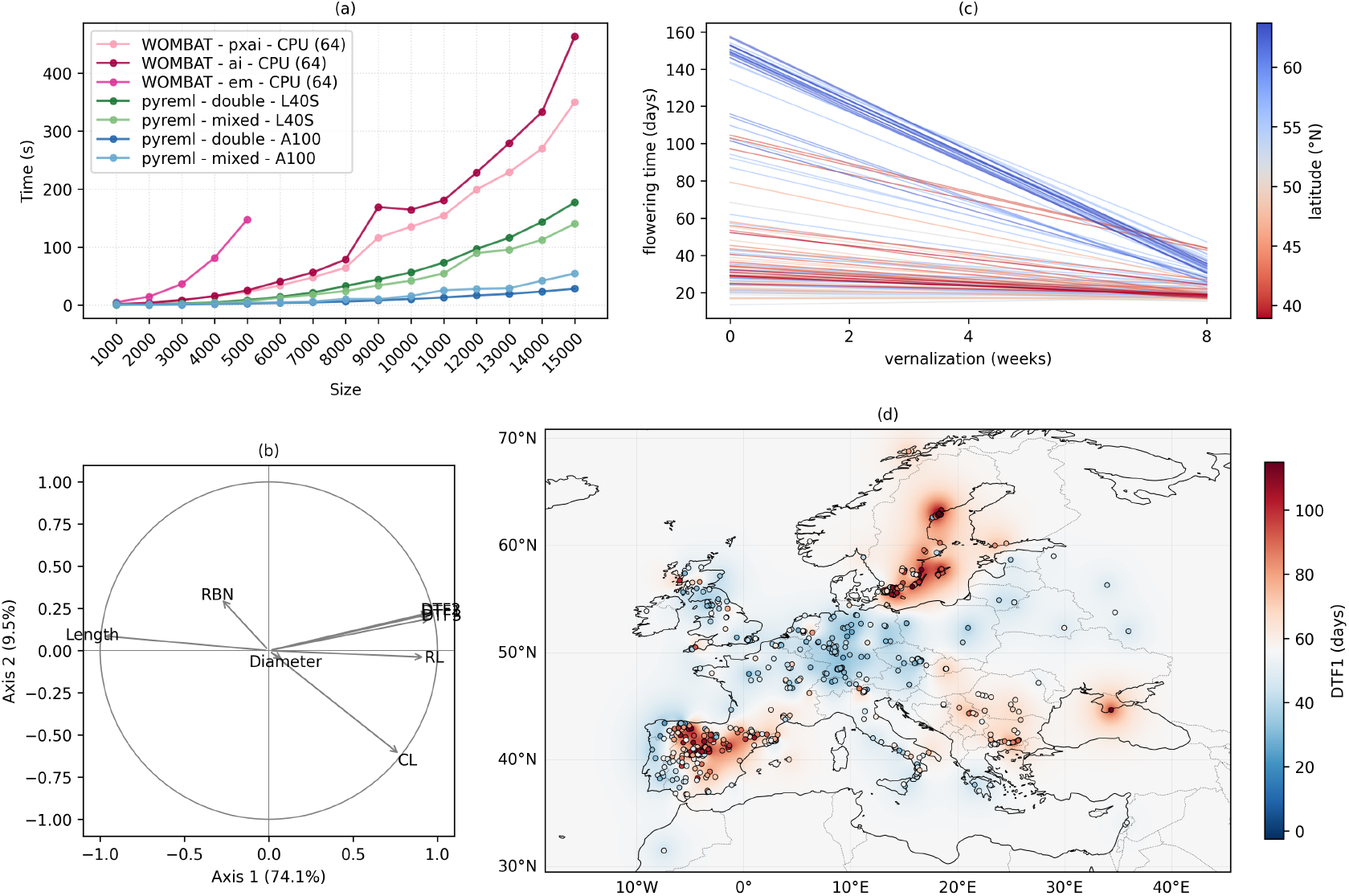
**(a)** Computation time depending on the software (`pyreml`, `Wombat`), its configuration (double numerical precision or mixed strategy for `pyreml`), and the hardware (CPU or GPU). **(b)** Correlation circle estimated using a factor analytic model between 8 phenotypic traits of *Arabidopsis thaliana*, centered and scaled to unit variance. **(c)** Analysis of the vernalization longitudinal data using random regression. **(d)** Spatial analysis of *Arabidopsis thaliana* days to flowering (DTF1).

We fitted a GBLUP model on an increasing sample size. A breeding-like population of *Arabidopsis thaliana* was simulated using `meiosim` (see Suppl. S3), a companion package that also ships all datasets used in this study. For each sample size *n*, the *n* × *n* kinship was built using a 5% minor allele frequency threshold, shrunk [Endelman and Jannink, 2012] and inverted before the timing started (see Suppl. S3.F1). All initial variances were taken to 1 for both `pyreml` and `Wombat`. We stopped running `Wombat` on EM algorithm for *n >* 5 000 as it was excessively slow.

All runs performed on `Wombat` or `pyreml` yielded the exact same variance components and accuracies, defined as cor(**u, û**), except for `Wombat` AI runs which did not converge for any *n* ≥ 10 000 (see Suppl. S4.F1). `pyreml` running on A100 was the fastest model of the benchmark, more than 12 times faster than `Wombat` on its best PX-AI algorithm, and more than 2 times faster when running on L40S (Fig. 1a).

The A100 GPU is optimized for double (FP64) operations, but most commercial GPUs are not. To maximize the benefit of GPU acceleration on all GPU types, we also implemented a mixed precision algorithm which consists in a warm-up on FP32 and polish on FP64. On FP64 GPUs (A100), the mixed approach slightly decreased performance, but it improved them in the more common FP32 GPUs (L40S) (Fig. 1a).

Finally, we pushed `pyreml` to an extreme GBLUP use case, analysing 25 000 simulated *A. thaliana* genotypes across 40 environments in a complete block design, that is one million observations. The intra-site heritability was lowered to a challenging but realistic 1%, and the 40 environments were both simulated and analysed as heteroscedastic. The analysis ran in 279 s in FP64 on the A100 GPU, and the resulting model reached an accuracy of 0.955. All variance parameters were well recovered, with a maximal relative error of 3.1%.

From the `pyreml` GBLUP model, a genome-wide association study (GWAS) can be straight-forwardly derived by backsolving [Gualdrón Duarte et al., 2014]. Backsolving from an LMM allows encompassing in a single model unbalanced observations, non-genotyped individuals [Wang et al., 2012], and including other effects such as non-additive genetic effects, environment and design effects, etc. Also, backsolving can follow a more complex GBLUP model such as a multivariate and/or longitudinal one.

### 3.2 pyreml is robust on challenging covariance structures

Factor Analytic (FA) primary goal is to reduce the number of variance parameters to estimate. However, it can also be seen as a direct inference on the latent structure of **Σ_a_** which is one of the most challenging problems a REML solver may address. Focus on the latent structure was exemplified by Marchal et al. [2026], from which we implemented most of `pyreml`’s FA algorithm. We benchmarked against `Wombat` again as it is the reference software for this purpose [Kirkpatrick and Meyer, 2004, Meyer, 2006, 2007, 2009].

We analyzed 8 phenotypic traits from the easyGWAS dataset [Grimm et al., 2017] of the 1001 genomes panel of *A. thaliana* [Alonso-Blanco et al., 2016], *with an order-2 FA model (****Σ*_*a*_***) and a kinship (****K*_*a*_***). The residual covariance was* ***R*** *=* ***Σ*_*R*_** *⊗* ***I*_*n*_** *with* ***Σ*_*R*_** *a dense, full rank 8 × 8 covariance between all traits. The initial variance structures were 0*.9 **I**_**p**_ + 0.1 **1_p_**×**_p_** for both **Σ**_**a**_ and **Σ_R_** in both `pyreml` and `Wombat`. As for the GBLUP benchmark, the kinship was built, shrunk and inverted before the timing started. The kinship **K_a_** was built from the 1001 genomes filtered on the RegMap panel (259 338 SNPs) [Horton et al., 2012, Pisupati et al., 2017]. After removing 2 individuals sharing more than 90% similarity with another retained individual to keep **K_a_** well-conditioned, we included *n* = 942 individuals with unbalanced observations per trait. `Wombat` was set to use the PX-AI algorithm and `pyreml` ran in double precision on the A100 GPU.

According to their own criterion, `pyreml` converged and `Wombat` did not. `pyreml` required a jitter *ε* = 10^−6^ on both the random and the residual variance structures for numerical stability. The estimates of the variance latent structure were similar between both software. It took 281 s for `Wombat` to yield these results, and 429 s (204 steps) for `pyreml` to converge. Loosening `pyreml` convergence criterion (Δ−2*l*_REML_) to meet the change in likelihood of the last `Wombat` step (Δ*l*_REML_ = 0.032223), `pyreml` yielded the same results in only 66 s (35 steps), confirming its greater computational speed (see Suppl. S4.F2).

The FA correlation circle presented Fig. 1b can be read as the one from a principal component analysis. The first axis shows the strong redundancy between the information carried by the phenotypic traits DTF1, DTF2, DTF3, length and RL. The second axis captures the remaining variance of CL, and the first factorial plan (axis 1 and 2 together) fails to describe both the diameter and the RBN.

FA models are especially meaningful in multi-environment trials, in which there is no residual correlation (the same experimental unit is never observed in different environment). In this case, the number of variance parameters grows linearly instead of quadratically with the number of responses, representing a meaningful gain in parsimony and computational cost. Though it has traditionally been limited to applications in quantitative genetics, the framework is agnostic and could be transposed to a much broader range of applications as suggested by Marchal et al. [2026].

### 3.3 Application to longitudinal data

One of the most common use of the mixed model is the analysis of repeated measurements, either controlled by quantitative covariates (random regression) or structured in groups (random factors). Applications range from merely getting rid of design effects (batches, blocs, confounding variables, etc.) to the study of growth curves, reaction norms, or factors of interest.

Still using the 1001 genomes filtered on the RegMap panel [Alonso-Blanco et al., 2016, Horton et al., 2012, Pisupati et al., 2017], we analyzed the *n* = 122 *A. thaliana* on which a vernalization experiment has been realized [Zhao et al., 2007, Atwell et al., 2010]. The experiment consisted in exposing the seeds to cold (*i*.*e*. vernalizing them) for 0, 2, 4 or 8 weeks, then measuring the number of days from sowing to flowering. Each phenotypic datapoint is an average over 10 plants for a given accession. The flowering date is right-censored at *y*_max_ = 125 days.

The model consisted in an order 1 (intercept and slope only) random regression of the flowering time over the number of vernalizing weeks. The random regression was structured with a kinship as described in the previous section. We compared the AIC for models with assumed homoscedastic residuals (model 0) versus heteroscedasticity learnt from the data, either as the form of a regression (**h** is an intercept and slope over the vernalizing weeks) (model 1) or as the form of a random factor (**h** is the vector of effect of each vernalization modality: 0 weeks, 2 weeks etc.) (model 2). Model 2 was better according to AIC (107.6 points lower than model 0 and 21.7 points lower than model 1) and we used it to predict the reaction norms of each accession. It ran in less than a second on the L40S GPU.

The reaction norms are presented Fig. 1c and colored relatively to the latitude of origin of the accessions. This visually confirms an expectable trend, which is that northern accessions seem more dependent on vernalization than southern ones. Further investigation could explore the incorporation of curvature using Legendre polynomials, as reviewed for instance by Schaeffer [2004].

### 3.4 Application to spatial analysis

Spatial analysis, which in a nutshell consists in estimating a variance structure and predicting new, unobserved points with kriging, is another one of the most standard LMM applications [Lark et al., 2006, Rousset and Ferdy, 2014]. We fitted a spatial model with an Euclidean decay kernel: (**K_a_**)_*ij*_ = exp(−*ρ* (**D**)_*ij*_), with **D** an assumed-Euclidean distance matrix.

We analyzed one-by-one 8 phenotypic traits of *A. thaliana*, grown in a common environment, depending on the coordinates of their origin [Alonso-Blanco et al., 2016, Grimm et al., 2017]. The analysis was restricted to individuals between 10^°^W and 40^°^E to retain a map mostly centered on Europe where the 1001 genomes origins are the most dense. For the trait DTF1, we ended up with *n* = 796 individuals and one observation per individual.

We compared `pyreml` to `spaMM` and found the former to be faster, although at this scale computation time is not a differentiating factor (about 6 seconds max). The outputs are strictly equivalent except for one edge case in which *ρ* is non identifiable (see Suppl. S4, Fig. S4.F3, Table S4.T1).

Using the out-of-the-box kriging tool implemented by `pyreml`, we predicted a grid of coordinates for the days-to-flowering 1 (DTF1) trait and presented it in Fig. 1d. The map displays the strong spatial structure of the day-to-flowering genetic determinism of *A. thaliana*.

As compared to specialized spatial analysis software, an LMM framework such as `pyreml` can combine in the same model a variety of fixed and random effects. The spatial component can be focal in the model, or it can be a potential confounding effect the user wants to account for in a broader model.

## 4 Discussion

Leveraging GPU acceleration, `pyreml` was unrivaled in the benchmarks that we presented. The computation times were especially low on A100, which is optimized for double precision operations. This illustrates a fundamental tension between the unavoidable accuracy required for LMM solving that can only be reached on double numerical precision, and the FP32 precision of most consumer GPUs.

`Wombat` is optimized for operations on sparse matrices, such as pedigree-based kinships [Meyer, 2006]. We confronted it on unfavorable ground, using a GBLUP model that requires dense matrix operations (Cholesky factorization, inversion). However, the genomic prediction era and the evolution towards the use of dense SNPs and whole genomes [Raimondi et al., 2022, 2024], may indicate that the next major challenges for mixed model application will lie in dense frameworks.

The core idea of `pyreml` is to solve REML by GD, which is a simple and generalist algorithm. It allows the use of a wide variety of variance structures, and will accommodate new needs and specific use cases that may arise. Furthermore, the low level model solver is abstracted from the high level API and `pyreml`’s engine may readily be repurposed to tackle atypical use cases requiring full control over **X, Z, G** and/or **R**, as for instance mixed effects neural networks [Xiong et al., 2019, Simchoni and Rosset, 2023].

Besides the computation speed and the internal design choices, `pyreml` provides a free, open source solution covering most of the functionalities a mixed model solver can provide. Generalized LMMs are not implemented, and could be considered for future evolution of `pyreml`. To our knowledge, no current open-source frequentist solver implements the factor-analytic mixed model, nor the most flexible heteroscedasticity specification from a mere formula. `pyreml` is available to all uses including commercial. Most importantly, the series of functional tests, as well as the present benchmark, demonstrated `pyreml` to be a reliable REML solver.

## Software availability

`pyreml` is distributed under a GPL3 license. It is available at: https://doi.org/10.5281/zenodo.21394319. The documentation is available at: https://ai-for-genome-interpretation-lab.github.io/pyreml/.

## Data availability

All datasets used in this study are distributed with the companion package `meiosim`, available at: https://doi.org/10.5281/zenodo.21134827.

