## Supplementary for "pyreml: a general-purpose, GPU-accelerated mixed model solver in Python"

#### S1 Documentation on training models

`pyreml` fits linear mixed models by direct differentiation of the Restricted Maximum Likelihood (REML) using `PyTorch` for variance parameter estimation. It benefits from `PyTorch` parallelization and GPU acceleration.

##### S1.1 Fitting process

###### S1.1.1 The `fit` method

Once a `pyreml` model is specified, it can be trained with the `.fit()` method.

```
model.fit()
```

This runs the full pipeline, calling this series of methods:

- `.OLS()`, Ordinary least square (OLS) initialization of $\beta$,
- `.REML()`, REML estimation of the variance components,
- `.HMME()`, Henderson's mixed model equations (HMME) for the random-effect predictions.

REML and HMME are engaged whenever the model carries random effects, a residual variance structure, or several responses; otherwise the fitting process stops at the OLS step.

###### S1.1.2 Numerical resolution

The `fit` method takes a `dtype` argument:

- `"mixed"` (default): OLS and a first REML pass run in `float32` to approach the optimum cheaply, then a second REML pass and HMME run in `float64` to refine the estimate and produce reliable outputs.

- `"double"`: all three steps (OLS, REML, HMME) run in `float64` (= `torch.double`). This option is intended for GPUs optimized for `float64` where the `float32` warm-up is not useful.
- `"float"`: all three steps run in `float32`. Fastest, but all output are single-precision and in some cases the REML solver can go off-track badly.

#### S1.2 OLS

The fixed effects are first estimated by ordinary least squares,

$$\hat{\beta}_{\text{OLS}} = (\mathbf{X}^\top \mathbf{X})^{-1} \mathbf{X}^\top \mathbf{y},$$

implying $\mathbf{X}$ to be full-rank. If the model carries no random effect, an `iid` residual and a single response, then the OLS solution is already the answer and training stops here. Otherwise, the OLS estimate initializes $\beta$ before REML. This initialization is always performed by `.fit()` and is strongly recommended for lower level uses of `pyreml`.

#### S1.3 REML

##### S1.3.1 REML loss

The per-effect blocks $\mathbf{G_a} = \mathbf{\Sigma_a} \otimes \mathbf{K_a}$ are assembled into the full random-effect covariance by a direct sum over effects:

$$\mathbf{G} = \bigoplus_{\mathbf{a}} \mathbf{G_a},$$

and the marginal covariance of the observations follows as:

$$\mathbf{V} = \mathbf{Z}\mathbf{G}\mathbf{Z}^\top + \mathbf{R},$$

Because observations may be missing in unbalanced designs, $\mathbf{\Sigma_R} \otimes \mathbf{K_R}$ is assembled over all original rows and then restricted to the observed entries through a residual “incidence” $\mathbf{W}$ such as: $\mathbf{R} = \mathbf{W}\,(\mathbf{\Sigma_R} \otimes \mathbf{K_R})\,\mathbf{W}^\top$.

The variance parameters and $\beta$ are estimated by direct gradient descent of the negative log-likelihood [Simchoni and Rosset, 2023], restricted (REML) as to yield unbiased estimates of the variance parameters [Patterson and Thompson, 1971, Harville, 1977]:

$$-2\ell_{\text{REML}} = \log|\mathbf{V}| + \mathbf{r}^\top \mathbf{V}^{-1} \mathbf{r} + \log|\mathbf{X}^\top \mathbf{V}^{-1} \mathbf{X}| + (n-p)\log(2\pi),$$

with $\mathbf{r} = \mathbf{y} - \mathbf{X}\beta$.

##### S1.3.2 Optimization

The objective is optimized with L-BFGS Liu and Nocedal [1989] under a strong-Wolfe line search, with Adam Kingma and Ba [2017] as a fallback for steps the line search cannot complete. The variance parameters and $\beta$ are optimized jointly.

##### S1.3.3 SMW

The REML computation uses the Sherman–Morrison–Woodbury identity (SMW) [Woodbury, 1950] which can have a very strong impact on the performance of the models depending on their specification.

**Interest of SMW for REML** The use of SMW aims at avoiding forming and inverting $\mathbf{V}$ directly:

$$\mathbf{V}^{-1} = \left(\mathbf{Z}\mathbf{G}\mathbf{Z}^\top + \mathbf{R}\right)^{-1} = \mathbf{R}^{-1} - \mathbf{R}^{-1}\mathbf{Z}\mathbf{C}^{-1}\mathbf{Z}^\top\mathbf{R}^{-1},$$

with the capacitance matrix $\mathbf{C}$:

$$\mathbf{C} = \mathbf{Z}^\top\mathbf{R}^{-1}\mathbf{Z} + \mathbf{G}^{-1}.$$

The associated determinant lemma gives the REML log-determinant without forming $\mathbf{V}$ either:

$$\log|\mathbf{V}| = \log|\mathbf{R}| + \log|\mathbf{G}| + \log|\mathbf{C}|.$$

Therefore, the problem transposes from inverting $\mathbf{V}$ to inverting $\mathbf{C}$, $\mathbf{G}$ and $\mathbf{R}$.

Using the properties of direct sum and Kronecker product:

$$\mathbf{G}^{-1} = \bigoplus_{\mathbf{a}} ({\mathbf{\Sigma}_\mathbf{a}}^{-1} \otimes {\mathbf{K}_\mathbf{a}}^{-1}).$$

For the residual:

$$\mathbf{R}^{-1} = \mathbf{W}\left({\mathbf{\Sigma}_\mathbf{R}}^{-1} \otimes {\mathbf{K}_\mathbf{R}}^{-1}\right)\mathbf{W}^\top,$$

which only stands under some conditions, including:

- if $\mathbf{W} = \mathbf{I}$;
- or if both $\mathbf{\Sigma}_\mathbf{R}$ and $\mathbf{K}_\mathbf{R}$ are diagonal, as $\mathbf{W}$ is a mere row-column selector.

Most left-hand and right-hand term inverses can be broken down even further, avoiding expensive computations for $\mathbf{G}^{-1}$ and $\mathbf{R}^{-1}$. The SMW path trades the complexity of inverting a dense $\mathbf{V}_{n\times n}$ for inverting a dense $\mathbf{C}_{q\times q}$ at each step.

**Implementation of SMW** SMW is usually more efficient when $q < n$, with $q$ the number of columns of $\mathbf{Z}$ (the total dimension of $\mathbf{u}$) and $n$ the number of stacked observations (*i.e.* $\mathbf{Z}$ is narrower than tall). Indeed, inverting the $q \times q$ matrix $\mathbf{C}$ is cheaper than inverting the $n \times n$ matrix $\mathbf{V}$ only when $q < n$. `pyreml` mostly conforms to this "dimensional rule", selecting `SMW=True` when $q < n$, `SMW=False` otherwise. However:

- using the high-level `from_dataframe` constructor, if the user forces `SMW=True` or `False` then their decision takes precedence over any other rule.
- using the high-level `from_dataframe` constructor, if all of the following conditions are met then SMW is switched off regardless of the dimensional rule:
  - $\mathbf{W} \neq \mathbf{I}$, which can arise for instance with several unbalanced options, or when using autoregressive structures.
  - the residual covariance $\mathbf{R}$ is not designed as a diagonal matrix.

Indeed, in this situation $\mathbf{R}^{-1} \neq \mathbf{W}\left({\mathbf{\Sigma}_\mathbf{R}}^{-1} \otimes {\mathbf{K}_\mathbf{R}}^{-1}\right)\mathbf{W}^\top$ and inverting the $n \times n$ matrix $\mathbf{R}$ is just as expensive as inverting $\mathbf{V}$.

- with the low-level constructor:
  - if `varmeth_inv` only is provided, SMW is forced on;
  - if `varmet` only is provided, SMW is forced off;
  - if both methods are available, the dimensional rule applies.

##### S1.3.4 Structured forward

Consider a random regression with three levels and four observations, an intercept and a covariate $x$ being carried on each level:

$$\mathbf{Z_a} = \begin{pmatrix} 1 & 0 & 0 & x_1 & 0 & 0 \\ 1 & 0 & 0 & x_2 & 0 & 0 \\ 0 & 1 & 0 & 0 & x_3 & 0 \\ 0 & 0 & 1 & 0 & 0 & x_4 \end{pmatrix}.$$

$\mathbf{Z_a}$ can be entirely described by the pair $(\mathbf{F_a}, s)$, defined as the values each observation carries and the level it loads them on:

$$\mathbf{F_a} = \begin{pmatrix} 1 & x_1 \\ 1 & x_2 \\ 1 & x_3 \\ 1 & x_4 \end{pmatrix}, \qquad s = (1, 1, 2, 3).$$

Row $i$ of $\mathbf{F_a}$ is written $\mathbf{f}_i$, and $s(i)$ is the level loaded by observation $i$. Over several responses, $\mathbf{F_a}$ is block-diagonal: each row carries the values of its own response, matching the response blocks of $\mathbf{\Sigma_a}$.

Assembling $\mathbf{V_a} = \mathbf{Z_a G_a Z_a}^\top$ directly forms the Kronecker block $\mathbf{G_a} = \mathbf{\Sigma_a} \otimes \mathbf{K_a}$, then multiplies through every column of $\mathbf{Z_a}$. Yet each row of $\mathbf{Z_a}$ being confined to a single level, entry $(i, j)$ of that product reduces to the covariance between the values carried by observations $i$ and $j$, scaled by the kernel entry between the levels they load:

$$[\mathbf{V_a}]_{ij} = \mathbf{f}_i^\top \, \mathbf{\Sigma_a} \, \mathbf{f}_j \, [\mathbf{K_a}]_{s(i)\, s(j)}.$$

The right-hand factor collects into $\mathbf{K_a}$ read at the observed level pairs:

$$\left[\mathbf{K_a}^{\text{obs}}\right]_{ij} = [\mathbf{K_a}]_{s(i)\, s(j)}, \qquad \mathbf{K_a}^{\text{obs}} = \begin{pmatrix} K_{11} & K_{11} & K_{12} & K_{13} \\ K_{11} & K_{11} & K_{12} & K_{13} \\ K_{21} & K_{21} & K_{22} & K_{23} \\ K_{31} & K_{31} & K_{32} & K_{33} \end{pmatrix},$$

so that the whole contribution is written as a Hadamard (elementwise) product $\odot$:

$$\mathbf{V_a} = \left(\mathbf{F_a} \, \mathbf{\Sigma_a} \, \mathbf{F_a}^\top\right) \odot \mathbf{K_a}^{\text{obs}}.$$

Neither $\mathbf{G_a}$ nor $\mathbf{Z_a G_a Z_a}^\top$ is ever formed, sparing both the memory and the time they would cost. This was exemplified with random regression, but holds for any effect built by `from_dataframe`. The full covariance follows as $\mathbf{V} = \sum_{\mathbf{a}} \mathbf{V_a}$, residual included.

On the SMW path (Suppl. S1.3.3), a second structured forward optimization is applied to prevent the formation of $\mathbf{R}^{-1}$ and of the Kronecker block $\mathbf{\Sigma_R}^{-1} \otimes \mathbf{K_R}^{-1}$ it is read from. Any $\mathbf{Z}$, $\mathbf{X}$ or $\mathbf{r}$ column can be reshaped as a matrix $\mathbf{U}$ over the level $\times$ response grid:

$$\mathbf{R}^{-1} \, \text{vec}(\mathbf{U}) = \text{vec}\left(\mathbf{K_R}^{-1} \, \mathbf{U} \, \mathbf{\Sigma_R}^{-1}\right).$$

The levels are contracted along the rows, the responses along the columns, and the result is flattened back. Two small products replace one large inverse, which is never formed. Unbalanced designs leave part of the grid empty, which is what the residual incidence $\mathbf{W}$ encodes. Operands are therefore lifted onto the complete grid, contracted, and read back at the observed cells afterwards.

The structured forward path is switched off when:

- using the `from_dataframe` high level builder, the user forces `structured_forward = False`;
- always when using the low level builder.

##### S1.3.5 Analytical backward

The REML gradient with respect to any variance parameter $\theta$ has a closed form:

$$\frac{\partial(-2\ell_{\text{REML}})}{\partial\theta} = \operatorname{tr}\left(\mathbf{A}\,\frac{\partial\mathbf{V}}{\partial\theta}\right),$$

$$\mathbf{A} = \mathbf{V}^{-1} - \mathbf{V}^{-1}\mathbf{r}\mathbf{r}^\top\mathbf{V}^{-1} - \mathbf{V}^{-1}\mathbf{X}\left(\mathbf{X}^\top\mathbf{V}^{-1}\mathbf{X}\right)^{-1}\mathbf{X}^\top\mathbf{V}^{-1},$$

with $\partial(-2\ell_{\text{REML}})/\partial\beta = -2\,\mathbf{X}^\top\mathbf{V}^{-1}\mathbf{r}$. As $\mathbf{V} = \sum_{\mathbf{a}}\mathbf{V}_{\mathbf{a}}$, for a given random effect (or the residual), the trace permutes into:

$$\frac{\partial(-2\ell_{\text{REML}})}{\partial\theta_{\mathbf{a}}} = \operatorname{tr}\left(\mathbf{Z}_{\mathbf{a}}{}^\top\mathbf{A}\,\mathbf{Z}_{\mathbf{a}}\,\frac{\partial\mathbf{G}_{\mathbf{a}}}{\partial\theta_{\mathbf{a}}}\right).$$

$\mathbf{A}$ is therefore evaluated once at the current point and shared by every variance parameter. Reusing the decomposition of Suppl. S1.3.4, where $\mathbf{Z}_{\mathbf{a}}$ is replaced by $(\mathbf{F}_{\mathbf{a}}, s)$ and $\mathbf{G}_{\mathbf{a}}$ by $\boldsymbol{\Sigma}_{\mathbf{a}}$ and $\mathbf{K}_{\mathbf{a}}$, the trace contracts onto each variance matrix:

$$\frac{\partial(-2\ell_{\text{REML}})}{\partial\boldsymbol{\Sigma}_{\mathbf{a}}} = \mathbf{F}_{\mathbf{a}}{}^\top\left(\mathbf{A}\odot\mathbf{K}_{\mathbf{a}}{}^{\text{obs}}\right)\mathbf{F}_{\mathbf{a}}, \qquad \left[\frac{\partial(-2\ell_{\text{REML}})}{\partial\mathbf{K}_{\mathbf{a}}}\right]_{st} = \sum_{\substack{i\,:\,s(i)=s\\ j\,:\,s(j)=t}}\left[\mathbf{A}\odot\mathbf{F}_{\mathbf{a}}\boldsymbol{\Sigma}_{\mathbf{a}}\mathbf{F}_{\mathbf{a}}{}^\top\right]_{ij}.$$

On the SMW path (Suppl. S1.3.3), $\mathbf{V}$ is never formed, so neither is $\mathbf{A}$. The trace is then read in the level space, from $\mathbf{Z}^\top\mathbf{A}\mathbf{Z}$ directly:

$$\mathbf{Z}^\top\mathbf{A}\mathbf{Z} = \mathbf{Z}^\top\mathbf{V}^{-1}\mathbf{Z} - \left(\mathbf{Z}^\top\mathbf{V}^{-1}\mathbf{r}\right)\left(\mathbf{Z}^\top\mathbf{V}^{-1}\mathbf{r}\right)^\top - \left(\mathbf{Z}^\top\mathbf{V}^{-1}\mathbf{X}\right)\left(\mathbf{X}^\top\mathbf{V}^{-1}\mathbf{X}\right)^{-1}\left(\mathbf{Z}^\top\mathbf{V}^{-1}\mathbf{X}\right)^\top$$

which uses three quantities the SMW projection already delivers through $\mathbf{C}$:

$$\mathbf{Z}^\top\mathbf{V}^{-1}\mathbf{M} = \mathbf{Z}^\top\mathbf{R}^{-1}\mathbf{M} - \left(\mathbf{Z}^\top\mathbf{R}^{-1}\mathbf{Z}\right)\mathbf{C}^{-1}\left(\mathbf{Z}^\top\mathbf{R}^{-1}\mathbf{M}\right), \qquad \mathbf{M}\in\{\mathbf{Z},\mathbf{r},\mathbf{X}\}$$

`PyTorch` autograd is therefore disabled through the assembly and factorisation of $\mathbf{V}$ ($\mathbf{C}$ on the SMW path). Automatic differentiation is only left with the last step, from each variance matrix to its own parameterization, and never traverses the Kronecker products.

The analytical backward path is switched off when:

- using the `from_dataframe` high level builder, the user forces `analytic_backward = False`;
- always when using the low level builder.

##### S1.3.6 Convergence criterion

Convergence is declared when two successive evaluations of $-2\ell_{\text{REML}}$ agree to within a tolerance $\tau$, the smallest difference in likelihood that remains meaningful given both the numerical resolution of the evaluation and the scale on which likelihoods are compared:

$$\tau = \min\left(\tau_{\max},\ |-2\ell_{\text{REML}}|\cdot 10^{-\delta_{\text{tot}}}\right), \qquad \delta_{\text{tot}} = \min\left(\delta_{\max},\ \delta_e - (\delta_c + \delta_r + \delta_\kappa)\right).$$

The restricted likelihood grows with the number of observations: the same dataset measured twice as often returns a value roughly twice as large. A fixed absolute threshold would therefore demand ever more significant digits as the dataset grows, and eventually more than the computer can hold. Counting significant digits keeps that demand constant. The budget starts from the machine epsilon $e$ of the working precision ($\delta_e = -\log_{10}(e) \approx 15.65$ in `float64`, $\approx 6.92$ in `float32`) and is capped by the conservative ceiling $\delta_{\max} = 10$. The three positive corrections count digits that are certain to be noise: $\delta_c$ carries the cumulated uncertainty of the elementary components the likelihood is summed from, while $\delta_r$ and $\delta_\kappa$ address the inversion of the variance $\mathbf{V}$ (or the capacitance $\mathbf{C}$ on the SMW path): $\delta_r$ as a random walk over the many operations of the factorisation, $\delta_\kappa$ as the information lost to the conditioning of the matrix. They are amplification factors that compound, hence the sum. Each is a conservative proxy, and $\delta_{\text{tot}}$ an estimate of the surviving resolution: no REML step is spent refining a decimal that carries no information.

The digit budget is relative, but what makes a likelihood difference meaningful is absolute: two models separated by $\Delta\text{AIC} = 2$ are considered distinguishable. The ceiling $\tau_{\max} = 0.02$, one hundredth of that scale, keeps the tolerance well below it. When the digit budget alone would push $\tau$ past $\tau_{\max}$, the noise floor of the evaluation has reached the scale at which models are told apart: the fit is reported as degenerate, the tolerance is clamped to $\tau_{\max}$, and convergence is denied.

The tolerance $\tau$ and the digit budget $\delta_{\text{tot}}$, $\delta_e$, $\delta_c$, $\delta_r$ and $\delta_\kappa$ are returned in the logs, where degenerate cases can be diagnosed.

##### S1.4 HMME

With the estimated variance components treated as known, the fixed and random effects are obtained jointly from Henderson's mixed model equations [Henderson, 1975]:

$$\begin{bmatrix} \mathbf{X}^\top\mathbf{R}^{-1}\mathbf{X} & \mathbf{X}^\top\mathbf{R}^{-1}\mathbf{Z} \\ \mathbf{Z}^\top\mathbf{R}^{-1}\mathbf{X} & \mathbf{Z}^\top\mathbf{R}^{-1}\mathbf{Z} + \mathbf{G}^{-1} \end{bmatrix} \begin{bmatrix} \hat{\beta} \\ \hat{\mathbf{u}} \end{bmatrix} = \begin{bmatrix} \mathbf{X}^\top\mathbf{R}^{-1}\mathbf{y} \\ \mathbf{Z}^\top\mathbf{R}^{-1}\mathbf{y} \end{bmatrix}.$$

Their solution gives the best linear unbiased estimator (BLUE) of $\beta$ and the best linear unbiased predictor (BLUP) of $\mathbf{u}$, in a single solve.

The same system also yields the error variances of the estimates and predictions.

##### S1.5 Akaike information criterion

Once the model is fitted, the Akaike information criterion (AIC) is computed in two different ways depending on whether it was performed by `OLS` or `HMME/REML`. The AIC is available as attribute `AIC` and the AIC method is consigned as `AIC_meth`.

These two AICs should not be compared one with each other. AIC computed post-OLS is designed for the exploration of fixed effects with no variance structure, and AIC computed post REML/HMME is designed for the investigation of the variance structure with a fixed $\mathbf{X}$ (*i.e.* a fixed `formula` in the model specification).

AIC post OLS is computed using the likelihood, from the unbiased residual variance $\sigma_R^2$:

$$-2\ell_{\text{ML}} = n \log\left(\frac{n-p}{n}\sigma_R^2\right) + n + n\log(2\pi)$$

and AIC post REML/HMME is computed using the restricted likelihood ($-2\ell_{\text{REML}}$).

In both cases, AIC is obtained by the addition of twice the total number of parameters ($\beta$, and all independent variance parameters).

#### S2 Functional tests

Every model is fitted on all combinations of the following flags:

- forcing the use and non-use of the SMW identity as both paths must return the same estimates, except for the autoregressive kernels which complete the full coordinate grid ($q > n$) and are never run with SMW.
- forcing the use and non-use of both the structured forward and analytical backward optimizations. When they are off, $\mathbf{V}$ (or the capacitance $\mathbf{C}$ under SMW) is assembled explicitly and the gradient is left to autograd.
- In the pedigree series the kernel is additionally supplied either as a covariance $\mathbf{K_a}$ or as an externally inverted precision matrix $\mathbf{K_a}^{-1}$ with its log-determinant $\log|\mathbf{K_a}|$. All tests run on CPU with the default mixed-precision strategy.

Most of these tests are based on the dataset `larix`, a subset of a *Larix* spp. genetic trial originally published with Marchal et al. [2017], limited to a few blocs of *Larix decidua* in one site, and distributed with `pyreml`. The interest of this dataset is to be both longitudinal, spatial, multivariate and genetically structured, making it a perfect sandbox for mixed modeling. Unless stated otherwise, the year-2000 campaign is used.

#### S2.1 Regression series

Height $y$ and circumference $x$ are centered and scaled. The fixed-only (OLS) model is fitted:

$$y_{ij} = \beta_0 + \beta_{bj} + \beta_1 x_{ij} + r_{ij}, \qquad r_{ij} \sim \mathcal{N}(0, \sigma_R^2),$$

with $\beta_0$ the intercept, $\beta_{bj}$ the effect of bloc $j$, and $\beta_1$ the effect of circumference of tree $ij$ on height (the slope). The outputs are compared to the `R` base function `lm`.

The random-regression model turns the bloc into a random intercept and slope:

$$y_{ij} = (\beta_0 + a_{0j}) + (\beta_1 + a_{1j})\, x_{ij} + r_{ij}, \qquad r_{ij} \sim \mathcal{N}(0, \sigma_R^2), \qquad \begin{bmatrix} \mathbf{a_0} \\ \mathbf{a_1} \end{bmatrix} \sim \mathcal{N}(\mathbf{0}, \mathbf{\Sigma_a} \otimes \mathbf{I}),$$

with $j$ the bloc and $ij$ the tree within it. It is compared to `lme4` [Bates et al., 2015] fitted with REML. All these tests pass.

#### S2.2 Spatial series

The series runs on a contrasted but compact bloc subset. Height is analyzed with a spatial model:

$$y_i = \mu + a_i + r_i, \qquad r_i \sim \mathcal{N}(0, \sigma_R^2), \qquad \mathbf{a} \sim \mathcal{N}(\mathbf{0}, \sigma_\mathbf{a}^2 \mathbf{K_a}),$$

with $a_i$ the effect of the coordinate of tree $i$ and $\mathbf{K_a}$ built from a spatial representation and learnt decay parameters. Five parameterizations of $\mathbf{K_a}$ are tested:

- a supplied covariance $\mathbf{K_a}$,
- a supplied distance matrix $\mathbf{D}$ from which $\mathbf{K_a}$ is computed as $\mathbf{K_a} = \exp(-\hat{\rho}\,(\mathbf{D})_{ij})$,
- a grid of coordinates $\mathbf{x}$ from which `pyreml` builds $\mathbf{D}$ using an Euclidean kernel $\mathbf{D} = \|\Delta\mathbf{x}\|$,
- an isotropic separable autoregressive kernel with one shared rate $\exp(-\hat{\rho}\sum_k |\Delta x_k|)$,
- an anisotropic autoregressive kernel $\exp(-\sum_k \hat{\rho}_k |\Delta x_k|)$.

References come from `rrBLUP` [Endelman, 2011], whose `mixed_solve` does not estimate decay rates: they are profiled outside the solver on the REML log-likelihood using `scipy.optimize`.

The anisotropic kernel is non-identifiable on this subset, where it absorbs enough of the field structure to drive $\sigma_R^2$ to zero. This leads to one documented `xfail` (duplicated with the structured forward / analytical backward flag) on the estimation of $\sigma_R^2$, all its other outputs being asserted. All other tests pass.

##### S2.3 Pedigree series

Pedigree based kinships **A** (additive) and **D** (dominance) are built by `pyreml` and checked against `nadiv` [Wolak, 2012], from which their algorithms is directly drown. Mixed model resolution is checked against `breedR` [Muñoz and Sanchez, 2026], which is a wrapper for `BLUPF90` [Misztal et al., 2014]. All models carry an intercept $\beta_0$ and a bloc effect $\beta_b$.

The univariate model encompasses 2 random effects:

$$y_{ij} = \beta_0 + \beta_{bj} + a_i + d_i + r_{ij}, \qquad r_{ij} \sim \mathcal{N}(0, \sigma_R^2),$$

with $a_i$ and $d_i$ the additive and dominance effects of tree $i$ in bloc $j$, such as $\mathbf{a} \sim \mathcal{N}(\mathbf{0}, \sigma_A^2\mathbf{A})$ and $\mathbf{d} \sim \mathcal{N}(\mathbf{0}, \sigma_D^2\mathbf{D})$. The two multivariate models analyze three traits (height, circumference and flexuosity) jointly:

$$y_{ijt} = \beta_{0t} + \beta_{bjt} + a_{it} + r_{ijt}, \qquad \mathbf{a} \sim \mathcal{N}(\mathbf{0}, \mathbf{\Sigma_a} \otimes \mathbf{A}), \qquad \mathbf{r} \sim \mathcal{N}(\mathbf{0}, \mathbf{\Sigma_R} \otimes \mathbf{I}),$$

with $t$ the trait, and $\mathbf{\Sigma_a}$, $\mathbf{\Sigma_R}$ either both diagonal or both "full" (unstructured). Among the many assertions being checked, the tests involve the prediction of new, unobserved individuals.

The full model carries two expected failures (`xfail`'s). Genetic and residual covariances trade against each other along a near-flat likelihood ridge, and under the SMW path the $\mathbf{\Sigma_a}$ assertion alone is exempted, its solution departing from `breedR`'s by less than 5%. The exemption is duplicated because of the covariance / precision flag as well as the structured forward / analytical backward flag. Noteworthily, this specification is demanding for `breedR` as well, whose average-information runs return a non-finite $-2\ell_{\text{REML}}$ on it. All other tests pass.

##### S2.4 Genomics series

This last series is different from the previous ones as, instead of comparing `pyreml` to another REML software, its absolute performance is asserted using simulation.

Marker genotypes are simulated from a `larix` pedigree, and a shrunk genomic relationship matrix $\mathbf{K_a}$ is computed and frozen [VanRaden, 2008, Endelman and Jannink, 2012]. `pyreml`'s kinship builder is asserted against `rrBLUP::A_mat`, from which its algorithm is drown. All other quantities against the simulation ground truth.

The first scenario shares one additive effect across environments with a heteroscedastic residual:

$$y_{ij} = \beta_0 + \beta_j + a_i + r_{ij}, \qquad \mathbf{a} \sim \mathcal{N}(\mathbf{0}, \sigma_a^2\mathbf{K_a}) \qquad r_{ij} \sim \mathcal{N}(0, \sigma_R^2 h_j),$$

with $i$ an individual tree, $j$ the environment, and $h_j$ the variance ratio between environment 0 and environment $j$ learnt from the data.

The second is multivariate over environments:

$$y_{ij} = \beta_{0j} + a_{ij} + r_{ij}, \qquad \mathbf{a} \sim \mathcal{N}(\mathbf{0}, \mathbf{\Sigma_a} \otimes \mathbf{K_a}), \qquad \mathbf{r} \sim \mathcal{N}(\mathbf{0}, \mathbf{\Sigma_R} \otimes \mathbf{I}),$$

with $\beta_{0j}$ an environment $j$ specific intercept, $a_{ij}$ the genetic effect of tree $i$ in environment $j$, $\mathbf{\Sigma_a}$ an order-2 factor-analytic covariance structure and $\mathbf{\Sigma_R}$ a diagonal variance matrix. Among other assertion checks, the latent structure that has been used for the simulation must be recovered.

The third is a bivariate random regression:

$$y_{tij} = (\beta_{0t} + a_{0ti}) + (\beta_{1t} + a_{1ti})\, x_{ij} + r_{tij}, \qquad \mathbf{a} \sim \mathcal{N}(\mathbf{0}, \boldsymbol{\Sigma}_{\mathbf{a}} \otimes \mathbf{K}_{\mathbf{a}}), \qquad \mathbf{r} \sim \mathcal{N}(\mathbf{0}, \boldsymbol{\Sigma}_{\mathbf{R}} \otimes \mathbf{I}),$$

with $\beta_{0t}$ the general intercept for trait $t$, $a_{0ti}$ the individual intercept modifier for individual $i$ on trait $t$, $\beta_{1t}$ the general slope for trait $t$, $a_{1ti}$ the individual slope modifier for individual $i$ on trait $t$, and $x_{ij}$, the $j^{\text{th}}$ observation of covariate $x$ for individual $i$.

The left-hand term $\boldsymbol{\Sigma}_{\mathbf{a}}$ is $4 \times 4$ over the response $\times$ term components. Four constrained structures are fitted: block-diagonal or proportional, blocked by response or by regression term. Assertions are structural as much as numerical: entries constrained to zero must be exactly zero, and proportional structures must show a constant block ratio equal to the reported coefficient.

All these tests pass.

### S3 Simulation

#### S3.1 How `meiosim` works

`meiosim` provides a single, lightweight data structure that bundles genotypes, individual metadata and a genetic map together. It includes the operations breeders and quantitative geneticists reach for most often:

- basic data manipulation,
- simulating meiosis and crosses,
- a ready-to-use *Arabidopsis thaliana* panel.

Everything in `meiosim` revolves around the `Population` object, which holds three aligned pieces of data:

- `genotypes`, an allele dosage coded as `0/1/2`,
- `metadata`, that guarantees three columns:
  - `individual`: a primary key for each individual,
  - `sire` / `dam`: parental identifiers, filled in by crossing operations.
- `map`, a marker map that holds the `chromosome` and `cM` columns.

And optionally:

- `phases`: a list `[hap0, hap1]` of two haplotype matrices, created by `phasing()` and propagated through crossing.
- `phenotypes`: a `pd.DataFrame` of trait values.

A shared `numpy` random generator (`rng`) makes every stochastic operation reproducible.

#### S3.2 Selection of operations

##### S3.2.1 `phasing()`

This method is strictly designed for fully inbred material whose phasing is trivial. For an actual phasing of heterozygous material an external tool must be used. Once the phases have been separated, they are indefinitely tracked in subsequent simulated crosses.

##### S3.2.2 `cross()`

Generate progeny from all pairwise combinations of the chosen parents. New simulated individuals inherit the right `sire` and `dam`, and a unique `individual` 16-character SHA-256 fingerprint.

##### S3.2.3 `selfing()`

Repeatedly self-fertilize one individual to drive it towards homozygosity, returning the population after `n_generations` rounds. The original parents are carried over into the resulting metadata.

#### S3.3 Arabidopsis dataset

The 1001 genome collection of *A. thaliana* [Alonso-Blanco et al., 2016] filtered on the RegMap panel [Horton et al., 2012, Pisupati et al., 2017] is provided as a resource.

The genetic map is built from the physical map assuming a constant 3.6 cM/Mb, simplifying the recombination landscape described by Salomé et al. [2012]).

Phenotypes come from Grimm et al. [2017] and Zhao et al. [2007, compiled by Atwell et al. 2010].

#### S3.4 Simulation scenario

The goal of the simulation is to produce a population with a breeding-like structure, *i.e.* a strong structuration in full- and half-sib families.

The mating design consisted in a diallel between 50 random founders sampled from the Swedish sub-population, crossed all together to produce 1 225 first generation hybrids (F1). From each F1, 100 lines are derived by 8 successive selfings yielding a strongly structured breeding population of 122 500 lines.

Twenty thousands (20 000) SNPs were sampled prior to the simulation, then split into 10 000 "simulated SNPs" and 10 000 "simulated QTLs". From the simulated QTLs, a single quantitative trait was simulated with an intercept of 100, and both a genetic and a residual variances of 10. The SNPs were reserved to build the kinship used for the GBLUP analysis (Figure S3.F1).

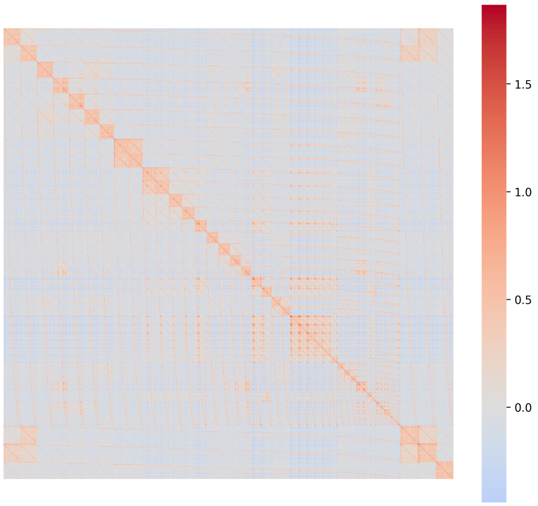


Figure S3.F1: Kinship built from the 10 000 SNPs of the 1 225 simulated F1 hybrids starting the *Arabidopsis* breeding population in a 50 × 50 diallelic mating design. Each F1 is then used to derive 100 fixed lines.

The analysis included an increasing number of individuals randomly sampled among the 122 500 simulated breeding lines.

### S4 Complementary results

#### S4.1 Upscaling GBLUP

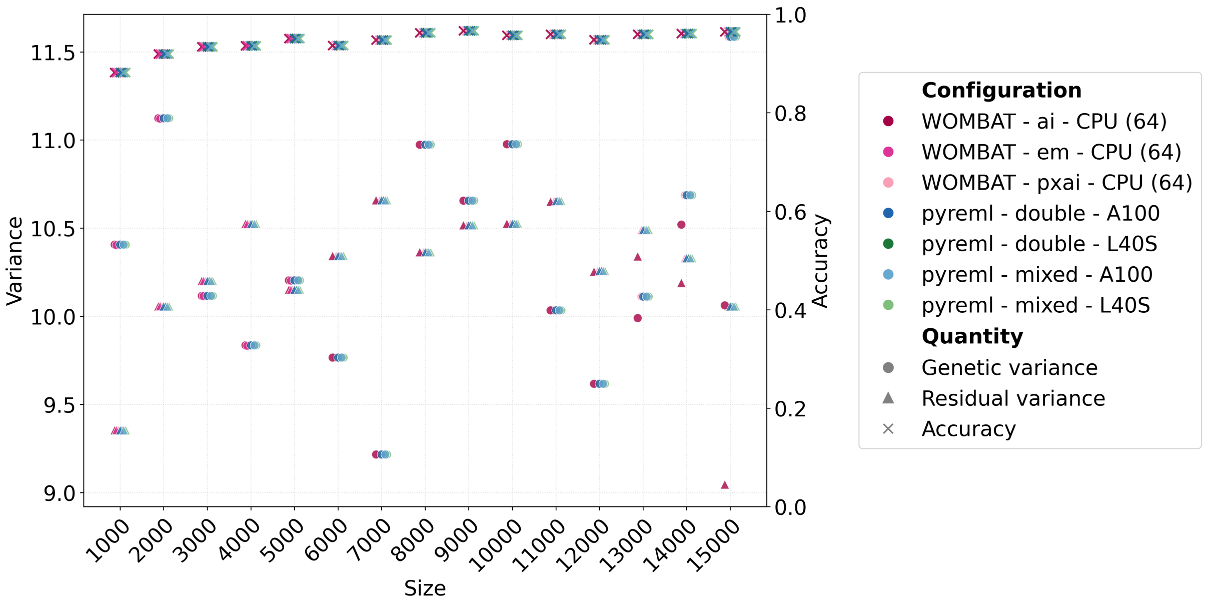


Figure S4.F1: Comparison of the model outputs depending on the model: variance estimation (left scale, simulated trait unit), and model accuracy defined as the correlation between predicted and simulated genetic values (right scale). Except for the `Wombat` average information (AI) runs that did not converge after size $\geq 10\,000$, all key outputs are strictly equivalent.

#### S4.2 FA model

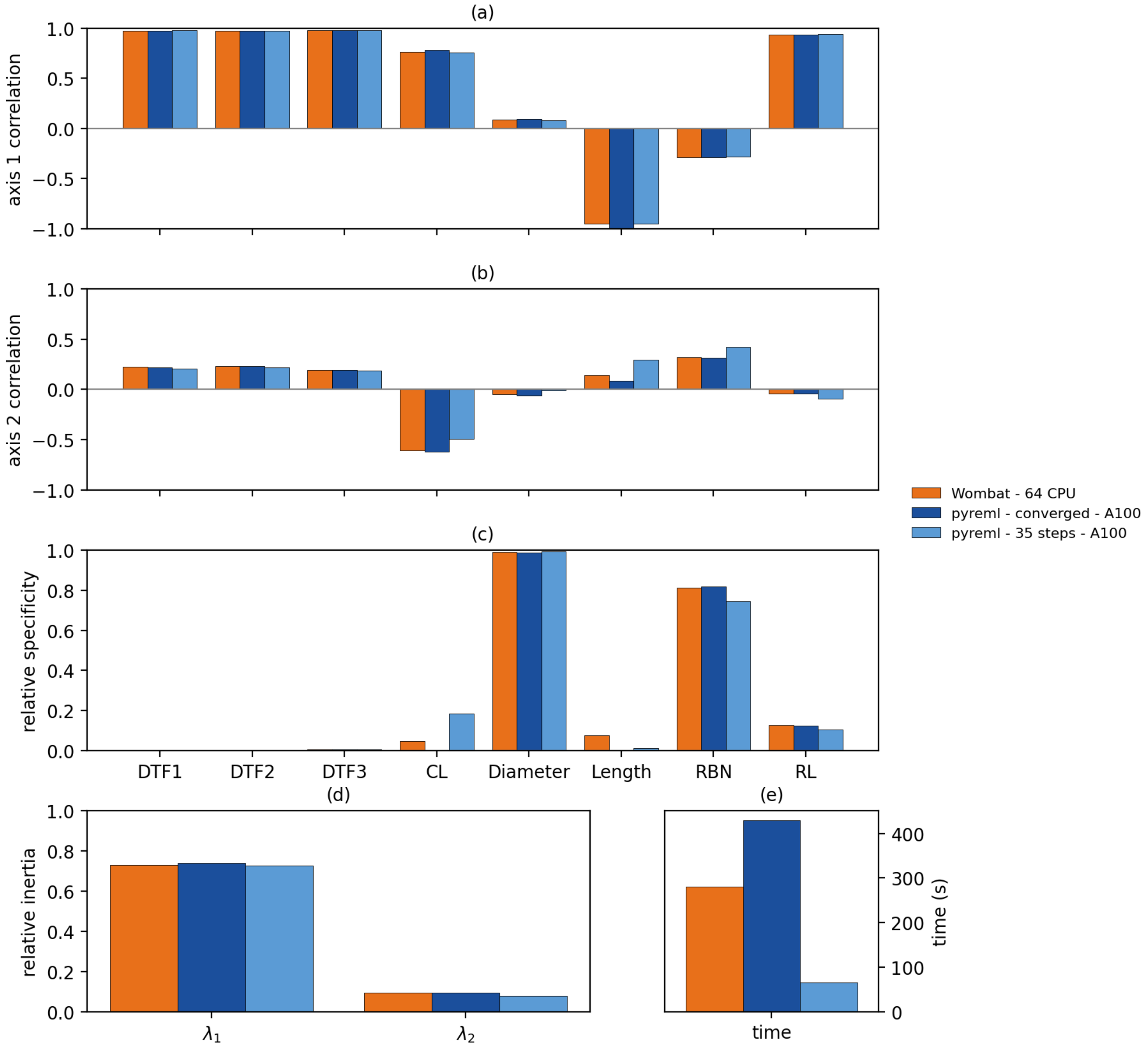


Figure S4.F2: Correlation to first (a) and second (b) factorial axis, (c) relative specificity defined as $\Psi_{\mathbf{a}\,ii}/\Sigma_{\mathbf{a}\,ii}$, (d) relative inertia defined as $\lambda_i/\text{trace}(\Sigma_{\mathbf{a}})$ and (e) computation time of the factor analytic model fitted with `pyreml` and `Wombat`. `Wombat` did not converge. However, all outputs are close except for the computation time.

#### S4.3 Spatial analysis

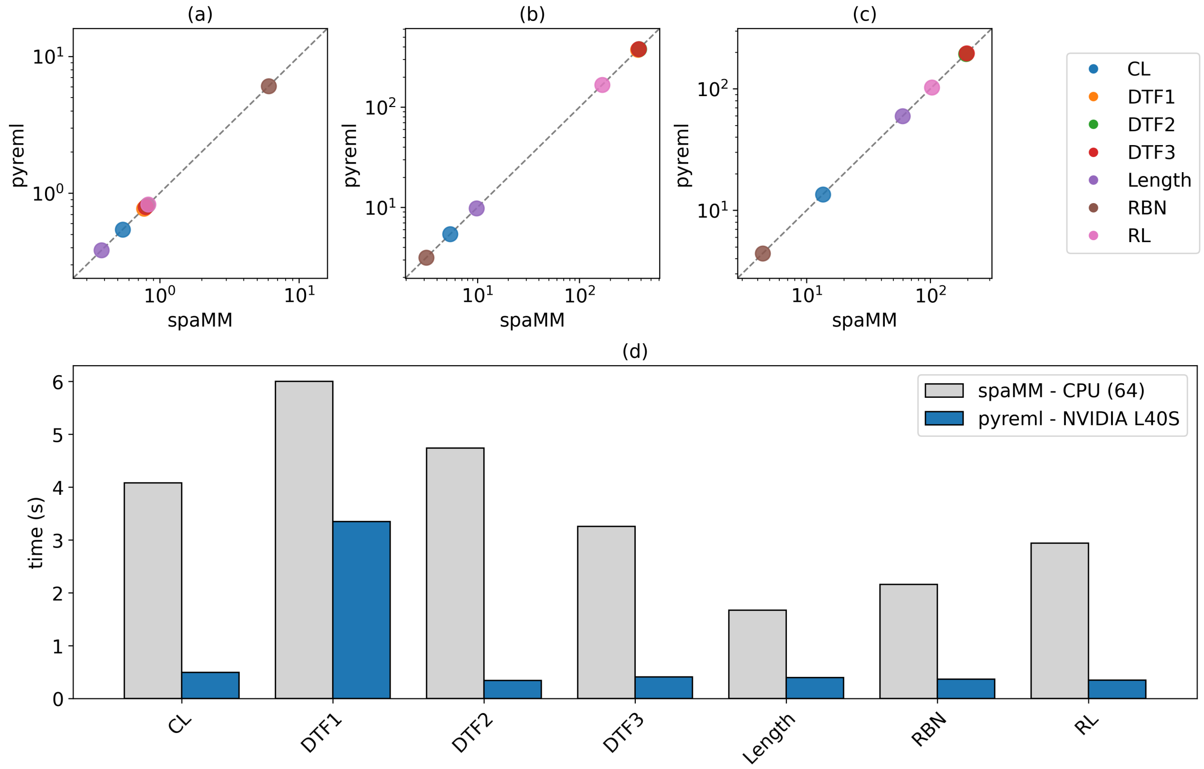


Figure S4.F3: Spatial analysis of seven phenotypic traits of Arabidopsis thaliana, comparing `pyreml` against `spaMM`. (a) Exponential decay parameter $\rho$, (b) spatial variance $\sigma^2_{\text{spatial}}$, (c) residual variance $\sigma^2_{\text{residual}}$, and (d) computation time for each phenotypic trait.

Table S4.T1: Estimation of the exponential decay $\rho$ and the variance parameters of spatial models, for each trait, using `pyreml` or `spaMM`. For the diameter, the two solutions lie on a flat ridge of the restricted likelihood leading to degenerate outputs. The REML likelihood was measured using `pyreml` in both cases.

| Solver | $\rho$ | $\sigma^2_{\text{spatial}}$ | $\sigma^2_{\text{residual}}$ | $-2\ell_{\text{REML}}$ |
|---|---|---|---|---|
| `pyreml` | 2.585e-08 | 5.523e+05 | 9.992 | 2865.623162 |
| `spaMM` | 0.002038 | 7.138 | 9.992 | 2865.657051 |
